# Social Motivation and Hippocampal-Cortical Structural Development in Adolescent Girls

**DOI:** 10.64898/2026.08.07.743556

**Authors:** Madeleine N. Goldberg, Ava J. Reck, Amalia M. Skyberg, Vishnu P. Murty, Jennifer H. Pfeifer

**Author notes:** Author Contacts: Madeleine N. Goldberg, Ava J. Reck, Amalia M. Skyberg, Vishnu P. Murty, Jennifer H. Pfeifer.

## Abstract

Adolescence is a fundamental developmental period marked by dramatic socio-affective and physiological changes, including shifts in social behavior and maturation of underlying brain architecture. In girls, this period coincides with the onset of the pubertal transition, which fundamentally influences motivated social behavior and neurodevelopment. The present study examines age- and pubertal maturation-related changes in social motivational goals and hippocampal and motivation-related cortical structural development in adolescent girls (*n*=154) across five timepoints. Social motivational goals showed substantial variability of each subdomain across age and pubertal development. Specifically, all social goals showed linear increases across age and pubertal stage, whereas goals centered around developing social competency increased non-linearly across age. Our neurodevelopmental findings align with established research, revealing volumetric increases of the hippocampus, and cortical thinning of the medial orbitofrontal cortex (mOFC) and rostral anterior cingulate cortex (rACC) across age and pubertal stages. Collectively, these results highlight simultaneous change in endorsement and prioritization of different social motivational goals across adolescence, and they underscore the simultaneous shifts in structural development in regions supporting social motivation and broader socio-affective development. This research highlights the importance of fostering positive social experiences during this critical developmental stage, with implications for adolescent well-being and social development.

## 1. Introduction

Adolescence is a crucial developmental period characterized by significant physical, neural, and socioemotional changes (Blakemore et al., 2010). Specifically, this period involves rapid maturation of brain structure (Vijayakumar et al., 2021) that is hypothesized to contribute to the development of reward-related cognitive functions such as decision-making, motivation, and risk-taking (Andrews et al., 2021; Chein et al., 2011). Additionally, adolescence is also a period of social and motivational changes, with peer relationships and experiences becoming increasingly salient and rewarding, particularly for adolescent girls (Pfeifer & Allen, 2021; Pfeifer & Berkman, 2018). The parallel timing of these changes has drawn significant attention to the pubertal transition, facilitated by the reactivation of the hypothalamic-pituitary-gonadal (HPG) axis (Naulé et al., 2021). The developmental processes underlying adolescent behavior are complex, requiring an integrated understanding of the neurobiological and social transformations that occur simultaneously during this critical period of development.

### 1.1 Social Development

The social landscape for young people transforms significantly during adolescence, with a clear shift in social motivations from being parent-focused to becoming increasingly peer-driven (Steinberg & Monahan, 2007). This transformation reflects behaviors aimed at “fitting in”, ranging from conformity to social norms to engagement in risky behaviors (Breiner et al., 2018; Helms et al., 2014; Neel et al., 2016). Additionally, peer influence during this time impacts adolescents’ navigation of social hierarchies, identity formation, and autonomy, which are crucial for their transition to adulthood (Blakemore & Mills, 2014; Crone & Dahl, 2012).

Social motivation shapes the evolution of prosociality across childhood, mirroring the social goals emerging during adolescence that drive peer engagement and provide the foundation for navigating complex social dynamics. Across adolescence, these goals evolve as they become more relevant to peer relationships and they reflect the increased neural sensitivity to social rewards. Prior work shows that agentic and communal social goals, reflecting egocentric and prosocial cognition, respectively, increase across early to mid-adolescence (Meisel et al., 2021; Trucco et al., 2014). These increases align with adolescents’ increased desire for social status and peer intimacy. Across adolescence, there are broad decreases submissive dimensions of these goals as desires for self-agency increase and salience of peer expectations decrease (Trucco et al., 2014). Further, the prioritization of these goals becomes more stable as youth progress through adolescence (Trucco et al., 2014).

There is also existing work that has disentangled how endorsement of agentic and communal social goals increases as a function of pubertal maturation in a sample of adolescents between 11 and 16 years old (Meisel et al., 2021). Specifically, within-subjects analyses show pubertal status is associated with higher levels of these social goals, and between-subjects analyses reflect this association in agentic goals only (Meisel et al., 2021). Further, girls endorse higher levels of communal social goals compared to boys (Meisel et al., 2021), which aligns with existing literature defining adolescence as a period of increased salience of girls’ peer relationships (Pfeifer & Berkman, 2018).

While limited to early and mid-adolescence, these studies lend insight into how social motivation changes across the adolescent social landscape. Both studies (Meisel et al., 2021; Trucco et al., 2014) use the *Interpersonal Goals Inventory for Children* (IGI-CR; Trucco et al., 2013), a self-report measure of agentic and communal goals, across three waves of data. Though a validated measure of social goals, the IGI-CR fails to capture goals that reflect the desire for creating and maintaining healthy peer relationships that do not necessarily include showing or avoiding social competence or incompetence, respectively (Scherrer et al., 2020).

Rudolph et al., 2011 categorizes social goals into three subdomains: 1) *social development goals*, focused on enhancing social skills and building sustainable, healthy peer relationships; 2) *demonstration-approach goals*, oriented toward gaining positive social recognition and prestige; and 3) *demonstration-avoidance goals*, aimed at minimizing negative judgments from peers. These dimensions offer valuable insight into adolescent social behavior, as they include proactive goals that require more nuanced cognitive reasoning beyond avoidance or approachfulness, while also reflecting both adolescents who may choose activities that align with their peers’ interests to gain approval, or avoid specific actions that could lead to social rejection.

The timing of this intensified sensitivity to peer feedback overlaps with normative neurodevelopmental changes, particularly in structures comprising motivational- and socioemotional-processing systems. The adolescent brain is especially adaptive to the heightened sensitivity to peer-related rewards and evolving social motivations, underscoring the role of the developing brain.

### 1.2 Adolescent neurodevelopment

The adolescent brain undergoes significant maturation in parallel with changes in social development, characterized by growth and notable reductions in brain volume and cortical thickness, respectively (Goddings et al., 2019). While there has been extensive research in exploring neural activity and development in reward-related and decision-making areas of the brain, this work has not been extended to the development of social motivation during adolescence.

Developmental research suggest that the ventromedial prefrontal cortex (vmPFC) plays a distinct role in assessing and integrating valence signals to update the value of potentially rewarding stimuli and outcomes (Blakemore & Robbins, 2012). Studies that have assessed the functional role of vmPFC during socially-rewarding contexts reveal that this region has a more distinct role in facilitating self-evaluation during adolescence (Barendse, Cosme, et al., 2020; Chavez et al., 2016; Pfeifer et al., 2013)and social comparisons between the self and similar peers in young adulthood (Moore et al., 2013). Additionally, improvements in cognitive control over value-based decisions during adolescence may be explained by a shift in projections from the vmPFC and other reward-related regions, to cognitive control areas such as the dorsolateral prefrontal cortex (Insel et al., 2019).

While these findings converge on the vmPFC as functionally relevant in socially-rewarding contexts, this region represents a broad anatomical boundary comprising multiple distinct subregions (Chase et al., 2020). Crucially, different parcellation approaches to capture mPFC may also include varying proportions of these subregions, each of which serves specialized functional roles (Chase et al., 2020). Additionally, focusing solely on reward-related regions in this way might overlook other regions may be more specialized and precise for social motivational processes.

Alternatively, one network that may be of interest when understanding adolescent social motivation is the mesolimbic system, where reward-related information is received by the medial orbitofrontal cortex (mOFC) and projected to the cingulate cortex and hippocampus (HPC) which prompt reward-guided decision-making and reward-related learning, respectively (Rolls, 2023). The mOFC is of particularly interest due to its role in ascribing value to prior feedback and experiences, helping to facilitate reward-guided decision-making (as discussed in Fettes et al., 2017), a cognitive process that is paramount to adolescent social development. Its projections to the cingulate cortex may also be relevant to social motivation, as the cingulate cortex is highly involved in self-evaluation and social comparisons, particularly when evaluating peers’ emotional states (Hamilton et al., 2012). All three regions undergo significant reductions in gray matter volume and cortical thickness across adolescence (Tamnes et al., 2017).

As previously mentioned, the HPC’s role in the mesolimbic system allows for rewarding information from the mOFC and cingulate cortex to be encoded and committed to episodic memory, providing information that helps set goals related to decision-making (Rolls, 2023). Further, this process may include the retrieval of socially-rewarding interactions, enhancing communication with other reward-related brain structures (see discussion in FeldmanHall et al., 2021). Prior developmental work has established this structure as crucial in guiding reward-motivation action, highlighting the dynamic interplay between memory systems and executive control during adolescence and into adulthood (Davidow et al., 2016; Elliott et al., 2022; Murty et al., 2016). Some research suggests that the growth of the HPC across critical periods of development (Lynch et al., 2019; Satterthwaite et al., 2013) may support social functioning and well-being (i.e., social transmission, social map-making; as discussed in Montagrin et al., 2018), pointing to a potentially broader role in this system supporting social motivation. A more integrative model that includes key mesolimbic structures and social goals may be essential to understand how teens navigate and prioritize social interactions and goals.

### 1.2 The present study

Social motivation has not been studied longitudinally; we don’t have an understanding on how specific dimensions of social motivation evolve across adolescence; vmPFC is too broad; HPC is largely understudied as a key ROI in motivation; age is used as a proxy for development (obscuring changes related to normative pubertal development);

The present study simultaneously addresses these limitations by use of a coordinated, multimodal approach to characterize changes in social motivational goals and structural changes in mOFC, rACC, and HPC architecture across adolescent development. Using linear mixed effects models, we examine how subdomains of social motivational goals, mOFC and rACC cortical thickness, and HPC grey matter volume change across age and pubertal maturation. We expect to the following results from our work:

1. Increases across each social motivational goal subdomain across age and pubertal maturation, reflecting prioritization of peer engagement during adolescence.
2. Non-linear reductions in cortical thickness of mOFC and rACC across age and pubertal maturation.
3. Non-linear increases in hippocampal grey matter volume across age and pubertal maturation.

By addressing each aim, our research seeks to deepen our understanding of how facets of social motivation are prioritized across adolescence, and how normative structural changes to brain architecture may support these socioemotional shifts.

## 2. Methods

### 2.1. Data Collection

The current study uses data from the Transitions in Adolescent Girls (TAG) project (see Barendse et al., 2020), an ongoing longitudinal study examining adolescent social and neural processes and mental health as a function of puberty-related changes. Here, 174 participants ages 10-13 years old were recruited from Lane County, Oregon from 2015-2018. Data was collected from participants in waves, initially spaced approximately 18 months apart, though interval lengths varied following the COVID-19 pandemic (Figure 1). Each visit was comprised of two sessions: Session 1 included (re)consent, a semi-structured diagnostic interview, and a batter of socioemotional questionnaires, and Session 2 entailed a thorough sequence of structural, resting-state, diffusion, and functional task MRI scans. Additionally, an in-lab hair sample for pubertal hormone assays, Oragene saliva kits for DNA collection, and anthropometric measure were acquired with another battery of questionnaires at Session 2. Between each session, participants complete up to 30 consecutive days of self-reports and provide weekly saliva samples across four weeks. The current study uses data collected from Waves 1 thorugh 5, and focuses socioemotional and pubertal development questionnaire, and structural MRI neuroimaging data.

**Figure 1.**
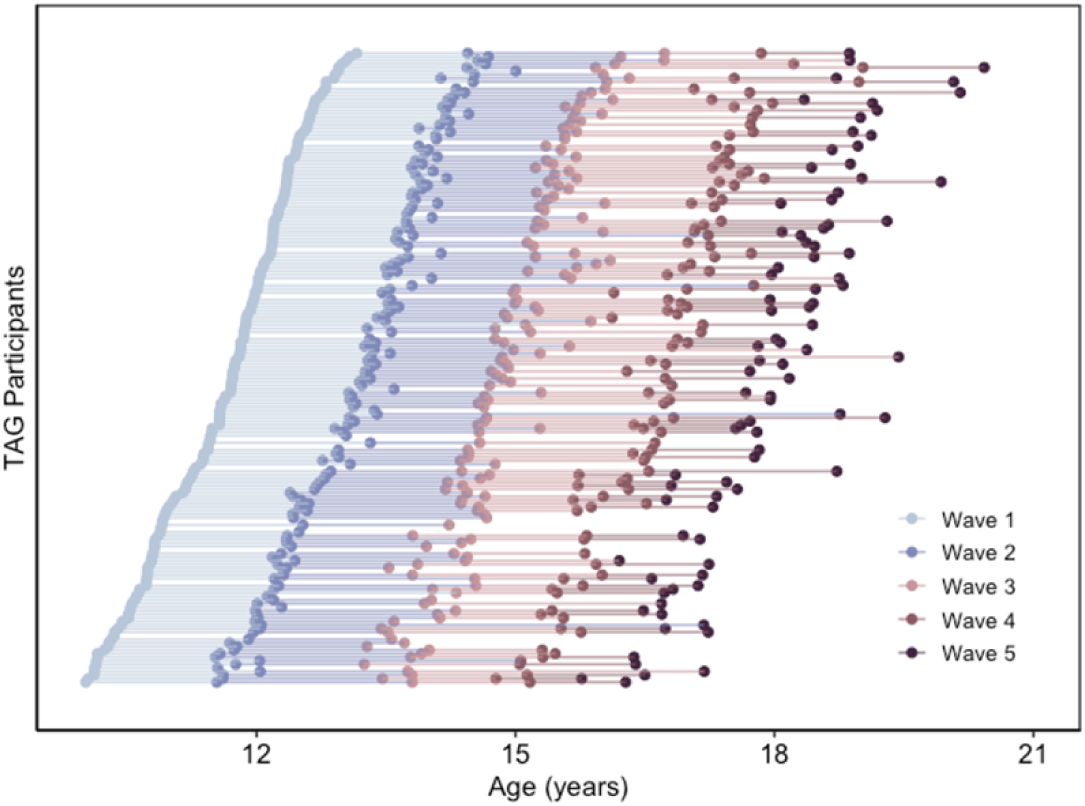
Participant Demographics. *Note.* Study participants across each timepoint. Each line represents participants’ age at each session across all timepoints.

### 2.2. Participants

At study onset, we included 174 participants aged 10-19 years (*M* = 11.68 years, *SD* = 0.80 years). All participants reported no existing history of significant medical or neurological disorders. Written informed consent and assent was obtained from participants along with parental consent for child participants.

### 2.3. Measures

#### 2.3.1. Structural MRI Procedures

Participants were screened for MRI safety compliance and practiced our larger study’s MRI protocol in a mock scanner so they would be comfortable in our magnet. Adolescents then completed a T1-weighted 3D MP-RAGE sequence (3.41ms TE; 2,500ms TR; 7 degree flip angle; 176 sagittal slices; 1mm isometric voxel size) in a 3T Siemens Skyra MRI scanner at the University of Oregon’s Lewis Center for Neuroimaging.

All T1-weighted structural scans were visually inspected for quality assurance before and after processing in FreeSurfer’s v8.0 longitudinal pipeline. Images were excluded from the current study’s analysis if they contained significant motion or susceptibility artifacts, excessive local noise, or fail reconstruction of our ROIs (mOFC, rACC, HPC). 20 scans were excluded in this process. The FreeSurfer longitudinal processing stream

For longitudinal analyses, all T1-weighted structural images were processed using the longitudinal stream in FreeSurfer v8.0 (Reuter et al., 2012). Each time point was first processed independently using the standard FreeSurfer recon-all pipeline, then an unbiased within-subject template was create from all available waves for each participant and used to guide the reprocessing of individual scans. This longitudinal approach uses the common template to initialize key processing steps, including skull stripping, Talairach registration, atlas registration, cortical surface reconstruction, and anatomical parcellation. By incorporating information across waves, the pipeline improves the reliability of estimates of cortical thickness, subcortical volume, and intracranial volume while reducing processing bias over time.

##### Social motivation

Social motivation was assessed at each timepoint using the Social Achievement Goals Questionnaire (SAQ; Rudolph et al., 2011). The SAQ was administered to participants at each timepoint and measuring three dimensions of social motivation: social development, which reflect a desire to develop social competence and positive relationships, demonstrative-approach goals, which reflect a desire to gain positive judgements from peers, and demonstrative-avoidance goals, which reflect a desire to avoid negative social judgements in peer interactions. The SAQ consists of 21 items that are rated on a 5-point Likert scale. Participants rate the extent to which each item describes their goals in peer interactions, with higher scores indicating greater endorsement of that social achievement goal orientation.

#### 2.3.2. Pubertal staging

Line Drawings (LD; Morris & Udry, 1980)) and the Pubertal Development Scale (PDS; Petersen et al., 1988)were used to measure a composite pubertal maturation measure. When completing the LD, participants identified which of the images of breast and pubic hair development best reflected their current developmental stage. Each stage reflects Tanner Staging, where development is scored on a 5-point scale ranging from 1 (*prepubertal*) to 5 (*complete maturation*). When completing the PDS, participants rated four questions about height, body hair, skin, and breast development on a 4-point scale ranging from 1 (*not started yet*) to 4 (*development seems complete*). An additional question related to menarche, or onset of menstruation, was presented as a yes/no item. To create a composite score of pubertal stage, the PDS scores were converted to Tanner Stages using a validated conversion syntax (Shirtcliff et al., 2009). The Tanner Stage scores from the LD and PDS were averaged to derive a composite pubertal maturation state (Byrne et al., 2023).

#### 2.3.3. Analysis plan

All statistical modeling analyses were conducted in R (https://www.r-project.org/). To account for any attrition across each wave of the study, we first assessed wither SAQ and sMRI data was missing completely at random (MCAR), assuming that the likelihood of missingness is unrelated to any potential confounding variables. Full-information likelihood estimation (FIML) was used to handle any missing data by testing hypotheses with all available data, yielding a more robust analysis despite any potential missing data.

All continuous outcome variables were winsorized to three standard deviations above and below their mean values to preserve the overall distribution of the data while reducing potential influence of outliers.

Linear mixed effects (LME) modeling was the optimal approach to fit the data due to its capacity to handle the complexities of repeated measures, nested data structures, and individual variability across multiple timepoints. LMEs also have the capacity to account for both fixed and random effects, a flexibility that is particularly ideal for analyzing longitudinal data with uneven intervals between observations and within-subject correlations (McCormick et al., 2023). Further, LMEs allow for the inclusion of covariates at both subject and timepoint levels, enabling more precise estimation of effects and a nuanced understanding of how predictors such as age or pubertal stage relate to behavioral and neural outcomes over time.

The present study considers linear, quadratic, and cubic LME models to fit our data. Linear models are the simplest approach to modeling a constant rate of change of a particular modality across a time-varying index of development (e.g., age or pubertal maturation). While these models suggest consistent, proportional changes, they fail to capture more complex, non-linear patterns in data. Quadratic models introduce a squared time-varying main effect term, producing a parabolic relationship, which may be useful when the data suggests a single inflection point.

To compare the fit of our LMEs under each condition and research question, we conducted ANOVA comparisons to evaluate model performance using likelihood ratio tests (LRT). LRTs allow us to determine whether additional complexity in models (e.g., additional parameters) significantly improve or hinder the fit to the data. Additionally, we used Akaike Information Criterion (AIC) and Bayesian Information Criterion (BIC) as additional metrics to assess model quality. AIC determines the goodness-of-fit of a particular model while penalizing any potential overfitting, and BIC favors simpler models. In general, lower AIC and BIC values indicate a better model fit. If an LRT indicated no significant improvement in model fit (*p* > .05), we selected the linear model.

## 3. Results

### SAQ Dimension Total Scores

Model comparisons using AIC indicated that the quadratic model provided the best fit for Social Developmental Goals across age (AIC = 3555.76), compared to the linear and cubic models. LRTs confirmed that quadratic term significantly improved fit over the linear models (*χ*^2^(1) = 6.69, *p* = .010), while the cubic term did not further improve fit over the quadratic model. Linear models provided the best fit for Approach and Avoidance Demonstration Goals across age. For Approach Demonstration Goals, the linear model had the lowest AIC (3246.36) relative to the quadratic and cubic models, and neither the quadratic nor cubic terms significantly improved fit. Similarly, for Avoidance Demonstration Goals, the linear model had the lowest AIC (3554.78) relative to the quadratic and cubic models, and neither the quadratic nor cubic terms significantly improved fit (Table 1, Figure 2).

**Figure 2.**
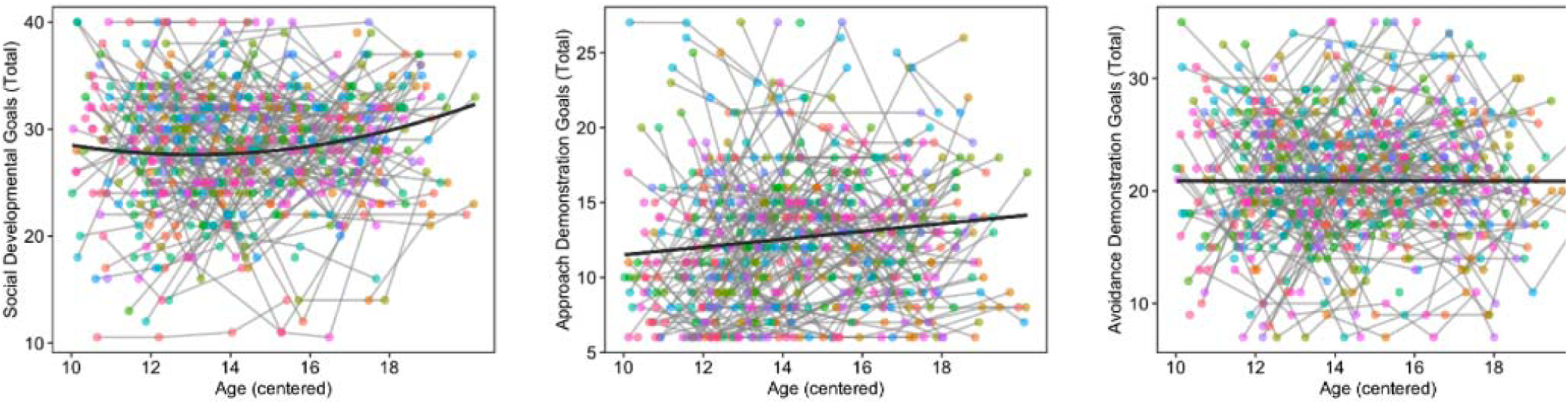
Social Motivational Goals by Age. *Note.* From left to right: Social Developmental Goals, Approach Demonstration Goals, and Avoidance Demonstration Goals. Colored points represent individual observations; gray lines connect repeated observations within participant across waves. Black lines represent model-predicted trajectories from the best-fitting linear mixed-effects model for each outcome. Age was mean-centered for model estimation (M = 14.38) and rescaled to raw values for display.

**Table 1.** Social Motivation Goals (Age)

|  | Model | df | AIC | BIC | logLik | Test | L.Ratio | p-value |
| --- | --- | --- | --- | --- | --- | --- | --- | --- |
| **Social Developmental Goals** |  |  |  |  |  |  |  |  |
| socdev_age_lin | 1 | 18 | 3560.45 | 3638.70 | -1762.22 |  |  |  |
| socdev_age_quad | 2 | 19 | 3555.76 | 3638.36 | -1758.88 | 1 vs 2 | 6.686 | 0.0097 |
| socdev_age_cubic | 3 | 20 | 3557.60 | 3644.55 | -1758.80 | 2 vs 3 | 0.162 | 0.6876 |
| **Approach Demonstrations** |  |  |  |  |  |  |  |  |
| app_age_lin | 1 | 18 | 3246.36 | 3324.52 | -1605.18 |  |  |  |
| app_age_quad | 2 | 19 | 3247.98 | 3330.48 | -1604.99 | 1 vs 2 | 0.384 | 0.5353 |
| app_age_cubic | 3 | 20 | 3249.37 | 3336.22 | -1604.69 | 2 vs 3 | 0.602 | 0.4380 |
| **Avoidance Demonstrations** |  |  |  |  |  |  |  |  |
| avoid_age_lin | 1 | 18 | 3554.78 | 3633.03 | -1759.39 |  |  |  |
| avoid_age_quad | 2 | 19 | 3556.73 | 3639.33 | -1759.37 | 1 vs 2 | 0.047 | 0.8280 |
| avoid_age_cubic | 3 | 20 | 3558.35 | 3645.30 | -1759.18 | 2 vs 3 | 0.377 | 0.5393 |

*Note.* Model comparisons were conducted using maximum likelihood (ML); final parameter estimates for the best-fitting model were obtained using restricted maximum likelihood (REML); *L.Ratio* and *p* reflect comparison to the preceding model in the table. AIC = Akaike Information Criterion; BIC = Bayesian Information Criterion.

Model comparisons across pubertal development indicated that linear models provided the best fit for all three SAQ dimensions. For Social Developmental Goals, although AIC was marginally lower for the quadratic model (AIC = 2921.60) than the linear model, this improvement did not reach statistical significance (χ²(1) = 2.67, *p* = .103), and the cubic term did not improve fit so the linear model was retained as the more parsimonious fit. For Approach Demonstration Goals, the linear model had the lowest AIC (2685.59) relative to the quadratic and cubic models, and neither the quadratic nor cubic term significantly improved fit. For Avoidance Demonstration Goals, the linear model similarly had the lowest AIC (2920.68) relative to the quadratic and cubic models, and neither the quadratic nor cubic terms significantly improved fit (Table 2, Figure 3).

**Figure 3.**
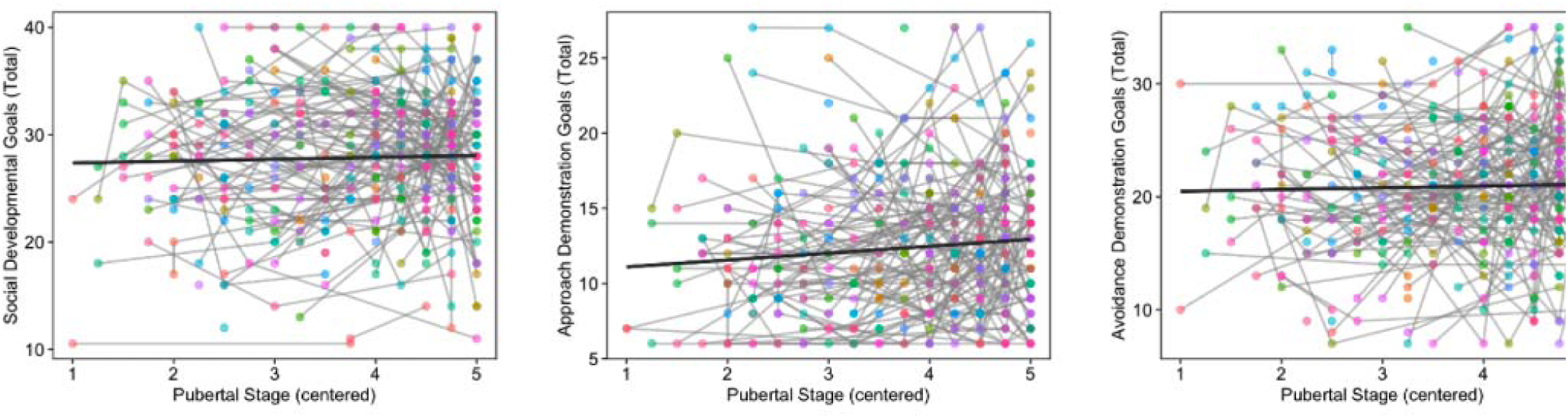
Social Motivational Goals by Pubertal Stage. *Note.* From left to right: Social Developmental Goals, Approach Demonstration Goals, and Avoidance Demonstration Goals. Colored points represent individual observations; gray lines connect repeated observations within participant across waves. Black lines represent model-predicted trajectories from the best-fitting linear mixed-effects model for each outcome. Pubertal stage was mean-centered for model estimation (M = 3.97) and rescaled to raw values for display.

**Table 2.** Social Motivation Goals (Puberty)

Social Motivation Goals (Puberty)
|  | Model | df | AIC | BIC | logLik | Test | L.Ratio | p-value |
| --- | --- | --- | --- | --- | --- | --- | --- | --- |
| **Social Developmental Goals** |  |  |  |  |  |  |  |  |
| socdev_pub_lin | 1 | 18 | 2922.27 | 2996.71 | -1443.13 |  |  |  |
| socdev_pub_quad | 2 | 19 | 2921.60 | 3000.18 | -1441.80 | 1 vs 2 | 2.666 | 0.1025 |
| socdev_pub_cubic | 3 | 20 | 2923.60 | 3006.31 | -1441.80 | 2 vs 3 | 0.000 | 0.9957 |
| **Approach Demonstrations** |  |  |  |  |  |  |  |  |
| app_pub_lin | 1 | 18 | 2685.16 | 2759.57 | -1324.58 |  |  |  |
| app_pub_quad | 2 | 19 | 2686.80 | 2765.34 | -1324.40 | 1 vs 2 | 0.361 | 0.5479 |
| app_pub_cubic | 3 | 20 | 2688.71 | 2771.38 | -1324.36 | 2 vs 3 | 0.090 | 0.7643 |
| **Avoidance Demonstrations** |  |  |  |  |  |  |  |  |
| avoid_pub_lin | 1 | 18 | 2920.68 | 2995.20 | -1442.34 |  |  |  |
| avoid_pub_quad | 2 | 19 | 2922.25 | 3000.90 | -1442.12 | 1 vs 2 | 0.434 | 0.5101 |
| avoid_pub_cubic | 3 | 20 | 2922.37 | 3005.17 | -1441.19 | 2 vs 3 | 1.874 | 0.1711 |

*Note.* Model comparisons were conducted using maximum likelihood (ML); final parameter estimates for the best-fitting model were obtained using restricted maximum likelihood (REML); *L.Ratio* and *p* reflect comparison to the preceding model in the table. AIC = Akaike Information Criterion; BIC = Bayesian Information Criterion.

### HPC GMV

Across age, model comparisons using AIC indicated that the cubic model provided the best fit for left hippocampal volume (AIC = 7284.83) relative to the quadratic and linear models, significantly improving model fit (χ²(1) = 25.15, *p* < .001). Similarly, the cubic model provided the best fit for right hippocampal volume across age (AIC = 7254.92) compared to the quadratic and linear models, significantly improving model fit (χ²(1) = 35.48, *p* < .001; Table 3, Figure 4).

**Figure 4.**
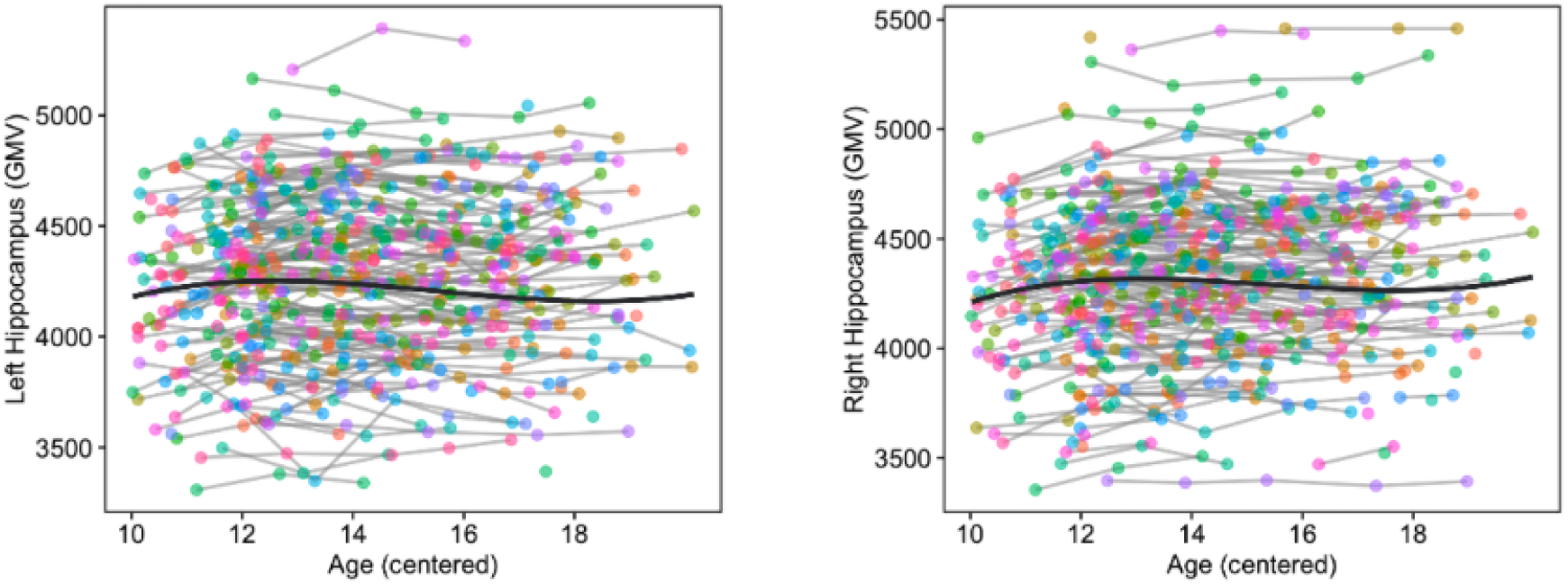
Hippocampus Volume by Age. *Note.* From left to right: Left Hippocampus volume, Right Hippocampus volume. Colored points represent individual observations; gray lines connect repeated observations within participant across waves. Black lines represent model-predicted trajectories from the best-fitting linear mixed-effects model for each outcome. Age was mean-centered for model estimation (M = 14.38) and rescaled to raw values for display.

**Table 3.** Hippocampus GMV (Age)

*Hippocampus GMV (Age)*
|  | Model | df | AIC | BIC | logLik | Test | L.Ratio | p-value |
| --- | --- | --- | --- | --- | --- | --- | --- | --- |
| **Left HPC** |  |  |  |  |  |  |  |  |
| lhpc_age_lin | 1 | 20 | 7311.55 | 7398.98 | -3635.77 |  |  |  |
| lhpc_age_quad | 2 | 21 | 7307.98 | 7399.79 | -3632.99 | 1 vs 2 | 5.563 | 0.0183 |
| lhpc_age_cubic | 3 | 22 | 7284.83 | 7381.01 | -3620.42 | 2 vs 3 | 25.152 | 0.0000 |
| **Right HPC** |  |  |  |  |  |  |  |  |
| rhpc_age_lin | 1 | 20 | 7291.75 | 7379.18 | -3625.87 |  |  |  |
| rhpc_age_quad | 2 | 21 | 7288.40 | 7380.21 | -3623.20 | 1 vs 2 | 5.343 | 0.0208 |
| rhpc_age_cubic | 3 | 22 | 7254.92 | 7351.10 | -3605.46 | 2 vs 3 | 35.484 | 0.0000 |

*Note.* Model comparisons were conducted using maximum likelihood (ML); final parameter estimates for the best-fitting model were obtained using restricted maximum likelihood (REML); *L.Ratio* and *p* reflect comparison to the preceding model in the table. AIC = Akaike Information Criterion; BIC = Bayesian Information Criterion.

Across pubertal maturation, model comparisons indicated that the cubic model also provided the best fit for left hippocampal volume (AIC = 6132.38) relative to the quadratic and linear models significantly improving model fit (χ²(1) = 5.56, *p* < .001). The quadratic model provided the best fit for right hippocampal volume across pubertal maturation (AIC = 6090.19) compared to the linear and cubic models, and neither linear nor cubic model significantly improved model fit (Table 4, Figure 5).

**Figure 5.**
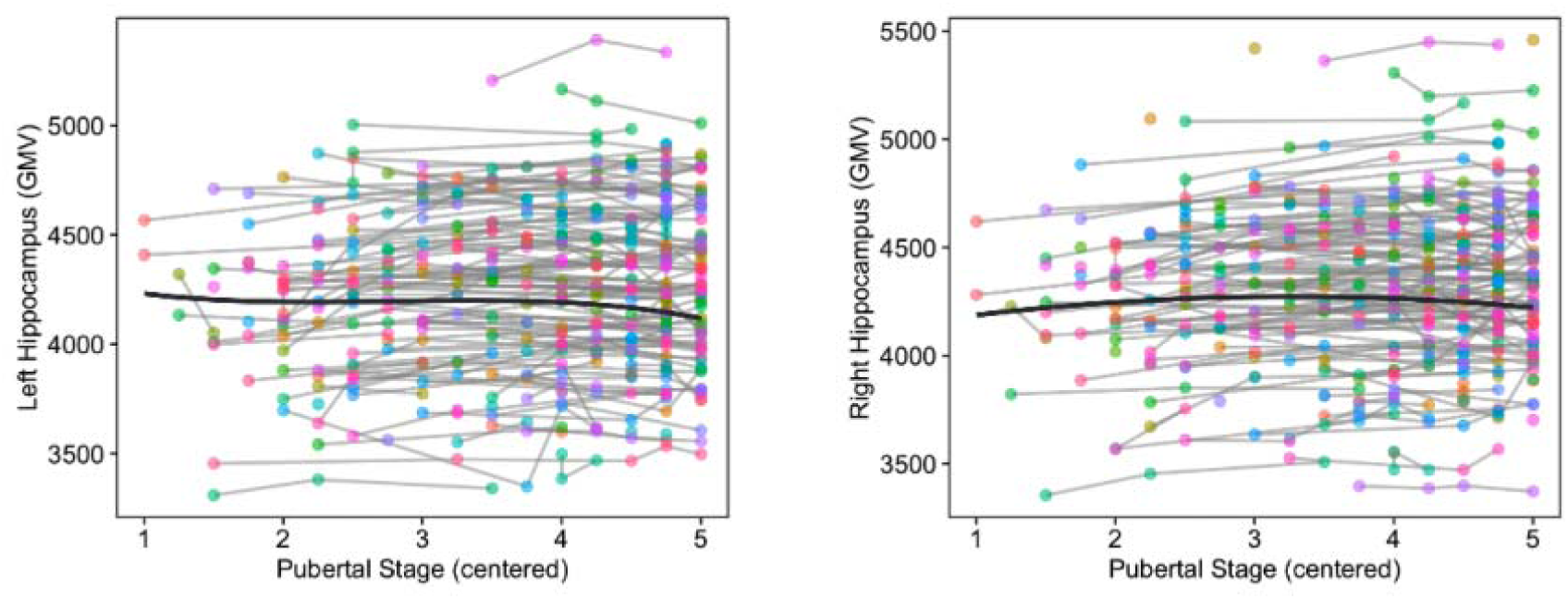
Hippocampus Volume by Pubertal Stage. *Note.* From left to right: Left Hippocampus volume, Right Hippocampus volume.

**Table 4.**
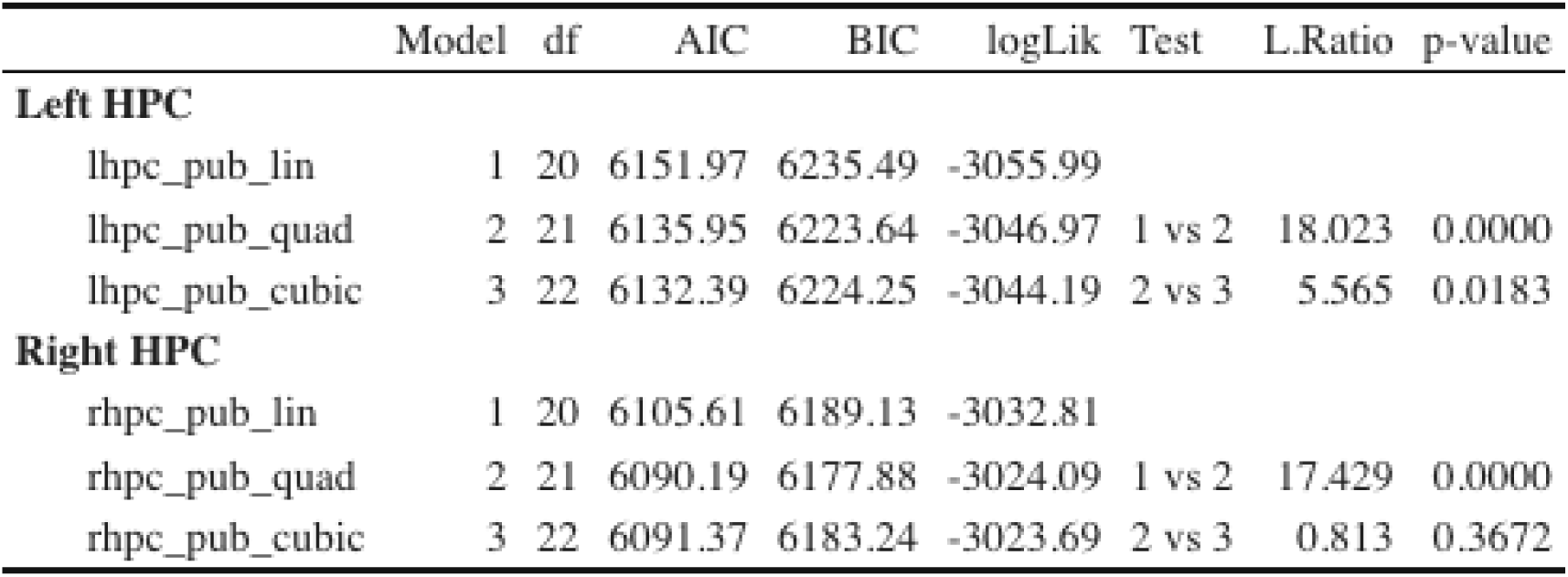
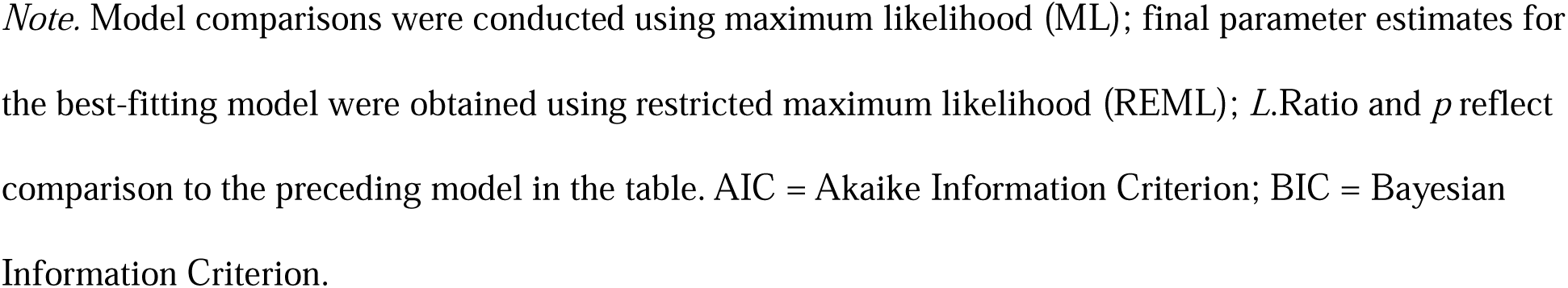
Hippocampus GMV (Puberty)

|  | Model | df | AIC | BIC | logLik | Test | L.Ratio | p-value |
| --- | --- | --- | --- | --- | --- | --- | --- | --- |
| **Left HPC** |  |  |  |  |  |  |  |  |
| lhpc_pub_lin | 1 | 20 | 6151.97 | 6235.49 | -3055.99 |  |  |  |
| lhpc_pub_quad | 2 | 21 | 6135.95 | 6223.64 | -3046.97 | 1 vs 2 | 18.023 | 0.0000 |
| lhpc_pub_cubic | 3 | 22 | 6132.39 | 6224.25 | -3044.19 | 2 vs 3 | 5.565 | 0.0183 |
| **Right HPC** |  |  |  |  |  |  |  |  |
| rhpc_pub_lin | 1 | 20 | 6105.61 | 6189.13 | -3032.81 |  |  |  |
| rhpc_pub_quad | 2 | 21 | 6090.19 | 6177.88 | -3024.09 | 1 vs 2 | 17.429 | 0.0000 |
| rhpc_pub_cubic | 3 | 22 | 6091.37 | 6183.24 | -3023.69 | 2 vs 3 | 0.813 | 0.3672 |

*Note.* Model comparisons were conducted using maximum likelihood (ML); final parameter estimates for the best-fitting model were obtained using restricted maximum likelihood (REML); *L*.Ratio and *p* reflect comparison to the preceding model in the table. AIC = Akaike Information Criterion; BIC = Bayesian Information Criterion.

### mOFC cortical thickness

Across age, model comparisons using AIC indicated that the linear model provided the best fit for left medial orbitofrontal cortical thickness (AIC = -1398.27) relative to the quadratic and linear models, with neither model significantly improving model fit. The cubic model provided the best fit for right medial orbitofrontal cortical thickness across age (AIC = -1302.47) compared to the quadratic and linear models, significantly improving model fit (χ²(1) = 13.33, *p* < .001; Table 5, Figure 6).

**Figure 6.**
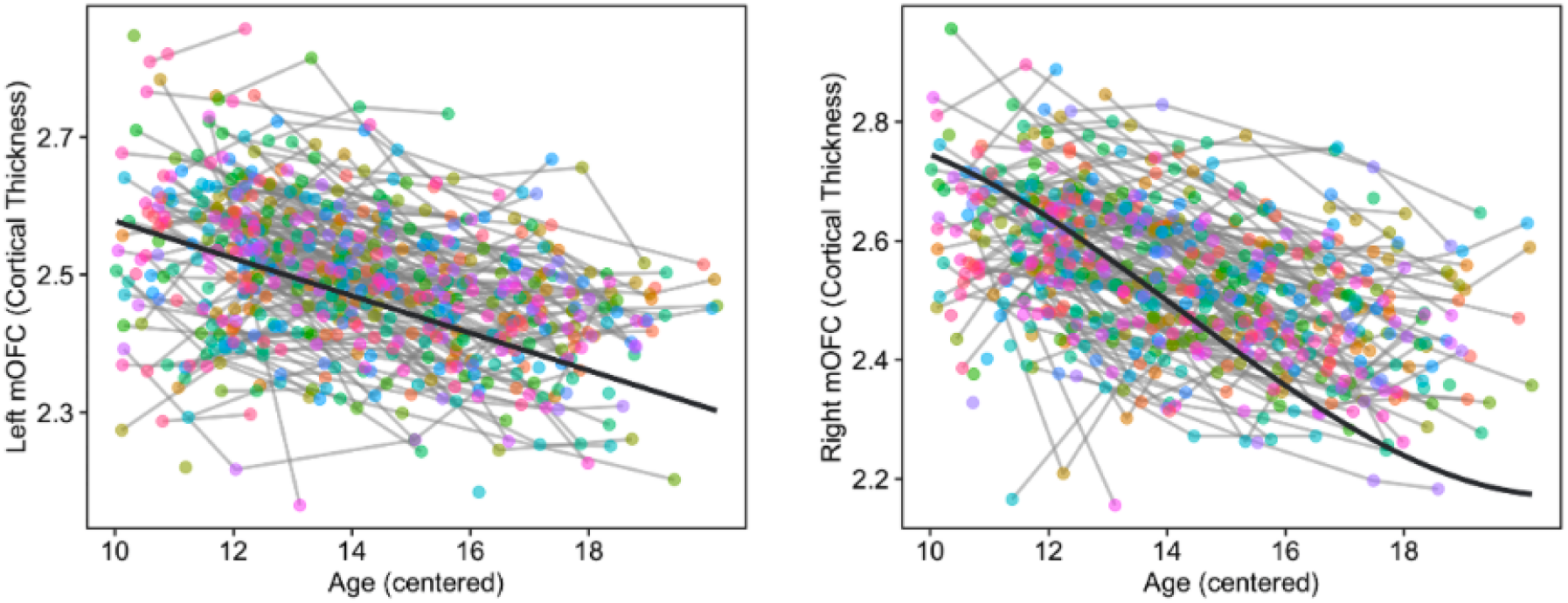
Medial Orbitofrontal Cortex Cortical Thickness by Age. *Note.* From left to right: Left Medial Orbitofrontal Cortex thickness, Right Medial Orbitofrontal Cortex thickness.

**Table 5.** Medial Orbitofrontal Cortex Cortical Thickness (Age)

Medial Orbitofrontal Cortex Cortical Thickness (Age)
|  | Model | df | AIC | BIC | logLik | Test | L.Ratio | p-value |
| --- | --- | --- | --- | --- | --- | --- | --- | --- |
| **Left mOFC** |  |  |  |  |  |  |  |  |
| lmoFc_age_lin | 1 | 20 | -1398.27 | -1310.84 | 719.134 |  |  |  |
| lmoFc_age_quad | 2 | 21 | -1396.68 | -1304.88 | 719.342 | 1 vs 2 | 0.417 | 0.5185 |
| lmoFc_age_cubic | 3 | 22 | -1398.00 | -1301.83 | 721.002 | 2 vs 3 | 3.320 | 0.0684 |
| **Right mOFC** |  |  |  |  |  |  |  |  |
| rmoFc_age_lin | 1 | 20 | -1292.35 | -1204.92 | 666.176 |  |  |  |
| rmoFc_age_quad | 2 | 21 | -1291.14 | -1199.34 | 666.570 | 1 vs 2 | 0.788 | 0.3749 |
| rmoFc_age_cubic | 3 | 22 | -1302.47 | -1206.29 | 673.235 | 2 vs 3 | 13.329 | 0.0003 |

*Note.* Model comparisons were conducted using maximum likelihood (ML); final parameter estimates for the best-fitting model were obtained using restricted maximum likelihood (REML); *L.Ratio* and *p* reflect comparison to the preceding model in the table. AIC = Akaike Information Criterion; BIC = Bayesian Information Criterion.

Across pubertal maturation, model comparisons indicated that the linear model provided the best fit for left medial orbitofrontal cortical thickness (AIC = -1109.41) relative to the quadratic and cubic models, with neither model significantly improving the model fit. The quadratic model provided the best fit for right medial orbitofrontal cortical thickness across pubertal maturation (AIC = -995.27) compared to the linear and cubic models, significantly improving model fit (χ²(1) = 15.34, *p* < .001; Table 6, Figure 7).

**Figure 7.**
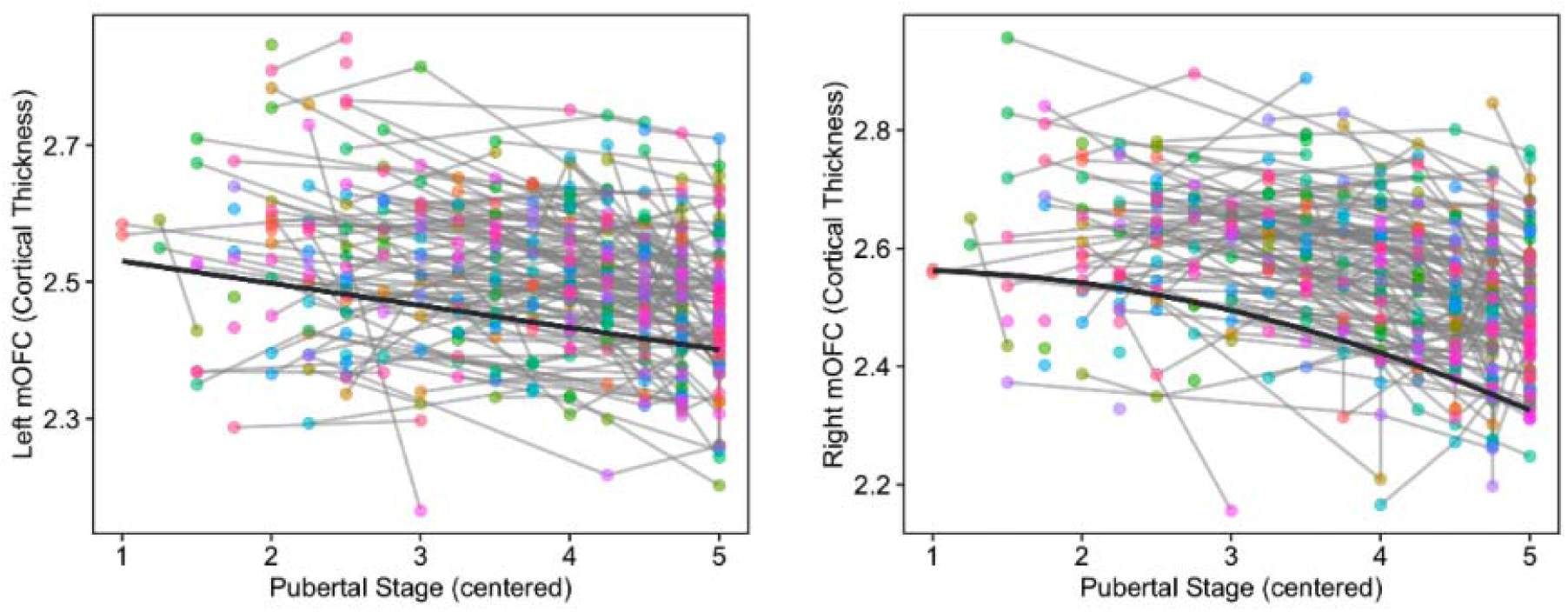
Medial Orbitofrontal Cortical Thickness by Pubertal Stage. *Note.* From left to right: Left Medial Orbitofrontal Cortex thickness, Right Medial Orbitofrontal Cortex thickness.

**Table 6.**
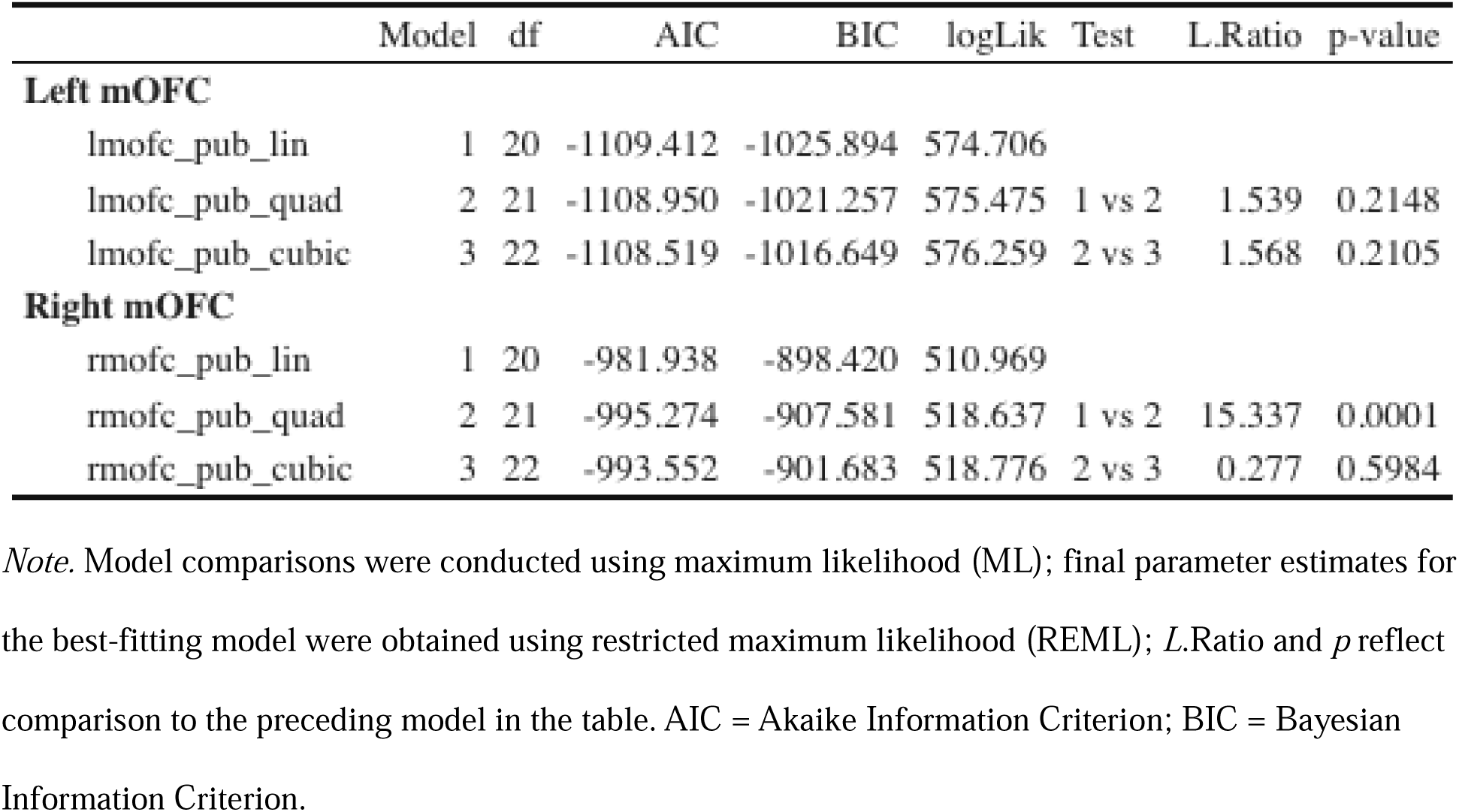
Medial Orbitofrontal Cortex Cortical Thickness (Puberty)

### rACC cortical thickness

Across age, model comparisons using AIC indicated that the cubic model provided the best fit for left rostral anterior cingulate cortical thickness (AIC = -1098.77) relative to the quadratic and linear models, significantly improving model fit (χ²(1) = 4.67, *p* = .030). Similarly, the cubic model provided the best fit for right rostral anterior cingulate cortical thickness across age (AIC = -1120.55) compared to the quadratic and linear models, significantly improving model fit (χ²(1) = 5.10, *p* = .023; Table 7, Figure 8).

**Figure 8.**
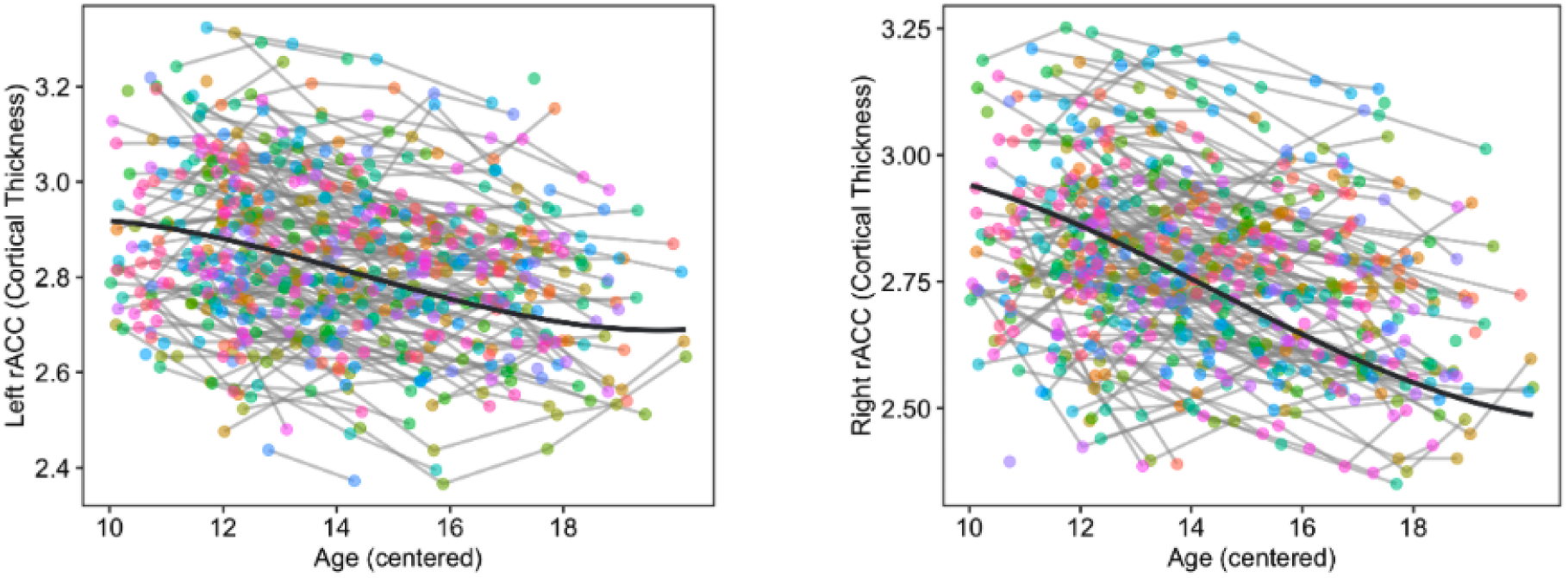
Rostral Anterior Cingulate Cortical Thickness by Age. *Note.* From left to right: Left Rostral Anterior Cingulate Cortex thickness, Right Rostral Anterior Cingulate Cortex thickness.

**Table 7.** Rostral Anterior Cingulate Cortex Cortical Thickness (Age)

Rostral Anterior Cingulate Cortex Cortical Thickness (Age)
|  | Model | df | AIC | BIC | logLik | Test | L.Ratio | p-value |
| --- | --- | --- | --- | --- | --- | --- | --- | --- |
| **Left rACC** |  |  |  |  |  |  |  |  |
| lracc_age_lin | 1 | 20 | -1098.10 | -1010.67 | 569.050 |  |  |  |
| lracc_age_quad | 2 | 21 | -1096.11 | -1004.30 | 569.054 | 1 vs 2 | 0.007 | 0.9348 |
| lracc_age_cubic | 3 | 22 | -1098.78 | -1002.60 | 571.387 | 2 vs 3 | 4.667 | 0.0307 |
| **Right rACC** |  |  |  |  |  |  |  |  |
| racc_age_lin | 1 | 20 | -1119.40 | -1031.97 | 579.703 |  |  |  |
| racc_age_quad | 2 | 21 | -1117.44 | -1025.64 | 579.721 | 1 vs 2 | 0.037 | 0.8471 |
| racc_age_cubic | 3 | 22 | -1120.55 | -1024.37 | 582.273 | 2 vs 3 | 5.104 | 0.0239 |

*Note.* Model comparisons were conducted using maximum likelihood (ML); final parameter estimates for the best-fitting model were obtained using restricted maximum likelihood (REML); *L.Ratio* and *p* reflect comparison to the preceding model in the table. AIC = Akaike Information Criterion; BIC = Bayesian Information Criterion.

Across pubertal maturation, model comparisons indicated that the linear model also provided the best fit for left rostral anterior cingulate cortical thickness (AIC = -819.97) relative to the quadratic and linear models, with neither model significantly improving model fit. For right rostral anterior cingulate cortical thickness across pubertal maturation, although the AIC was marginally lower for the cubic model (AIC = -844.69) than the quadratic and linear models, this improvement did not reach statistical significance (χ²(1) = 2.04, *p* = .153), and the quadratic model did not improve model fit (χ²(1) = 3.81, *p* = .051), so the linear model was retained as the more parsimonious fit (Table 8, Figure 9).

**Figure 9.**
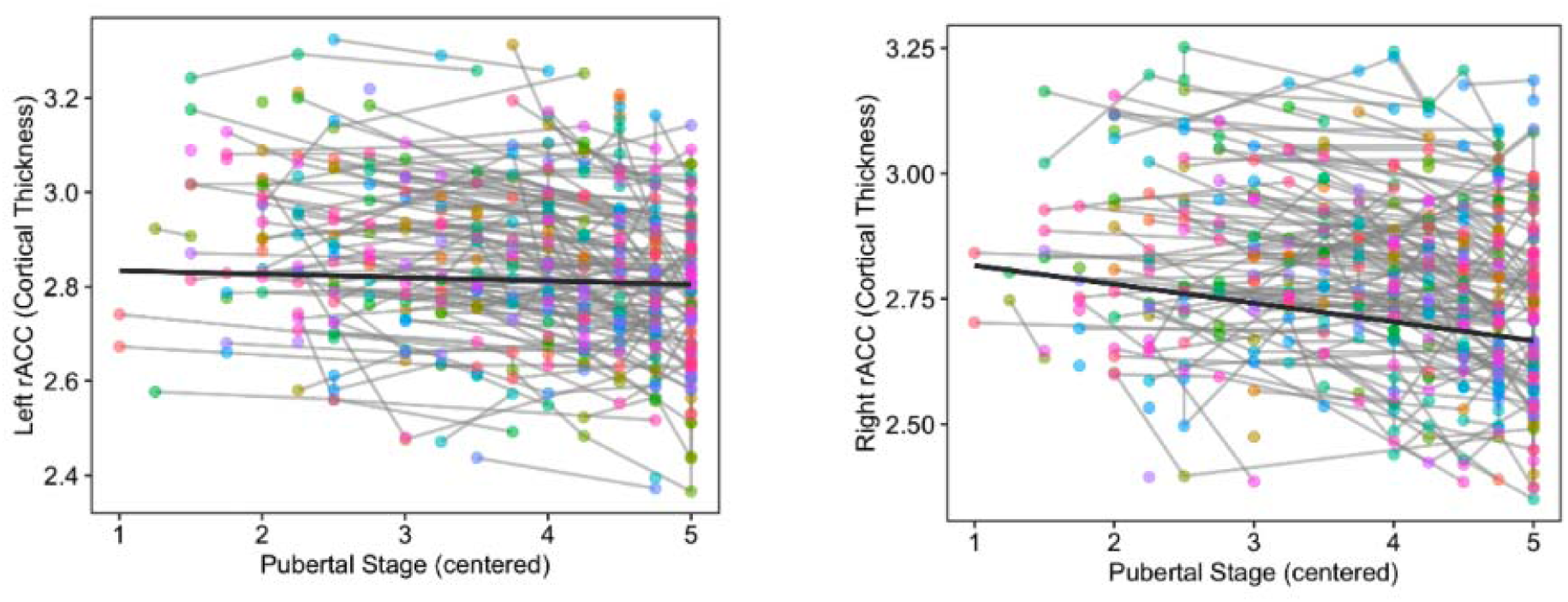
Rostral Anterior Cingulate Cortical Thickness by Pubertal Stage. *Note.* From left to right: Left Rostral Anterior Cingulate cortical thickness, Right Rostral Anterior Cingulate cortical thickness.

**Table 8.**
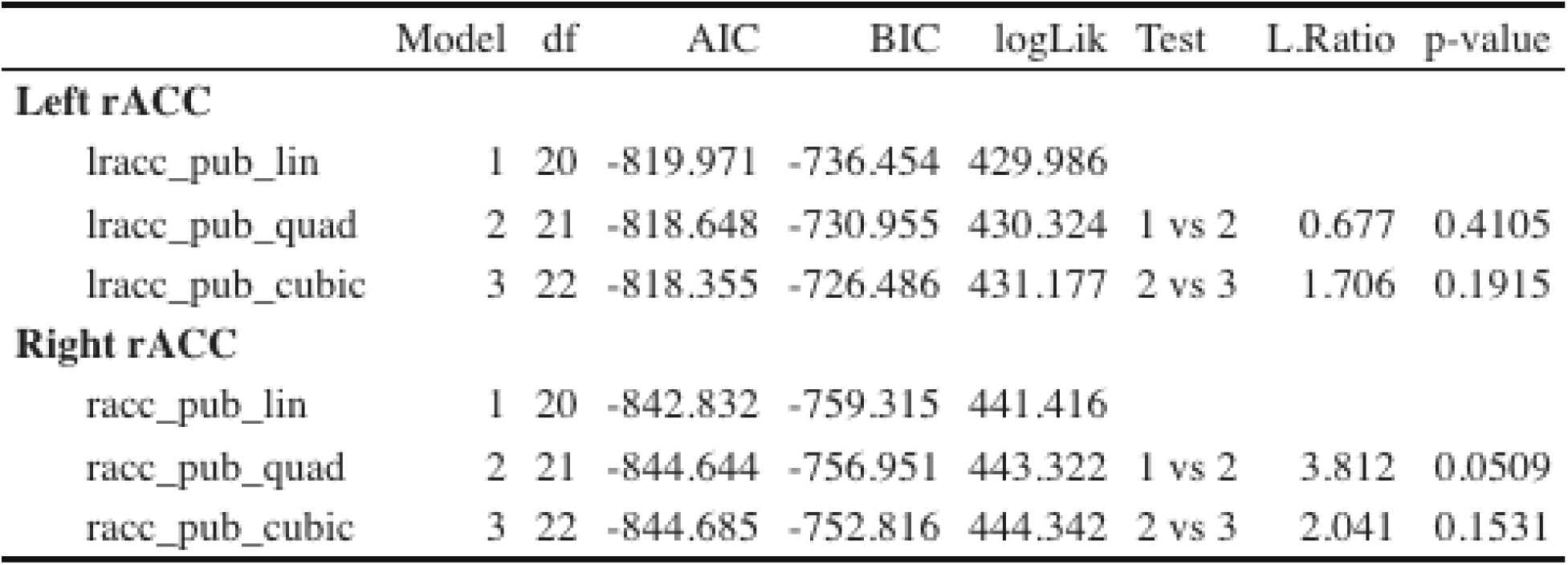
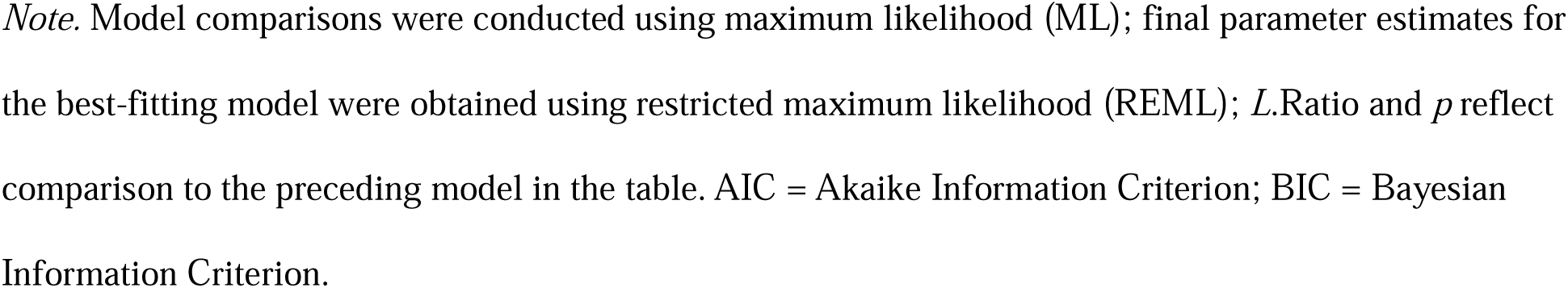
Rostral Anterior Cingulate Cortex Cortical Thickness (Puberty)

|  | Model | df | AIC | BIC | logLik | Test | L.Ratio | p-value |
| --- | --- | --- | --- | --- | --- | --- | --- | --- |
| **Left rACC** |  |  |  |  |  |  |  |  |
| lracc_pub_lin | 1 | 20 | -819.971 | -736.454 | 429.986 |  |  |  |
| lracc_pub_quad | 2 | 21 | -818.648 | -730.955 | 430.324 | 1 vs 2 | 0.677 | 0.4105 |
| lracc_pub_cubic | 3 | 22 | -818.355 | -726.486 | 431.177 | 2 vs 3 | 1.706 | 0.1915 |
| **Right rACC** |  |  |  |  |  |  |  |  |
| racc_pub_lin | 1 | 20 | -842.832 | -759.315 | 441.416 |  |  |  |
| racc_pub_quad | 2 | 21 | -844.644 | -756.951 | 443.322 | 1 vs 2 | 3.812 | 0.0509 |
| racc_pub_cubic | 3 | 22 | -844.685 | -752.816 | 444.342 | 2 vs 3 | 2.041 | 0.1531 |

*Note.* Model comparisons were conducted using maximum likelihood (ML); final parameter estimates for the best-fitting model were obtained using restricted maximum likelihood (REML); *L*.Ratio and *p* reflect comparison to the preceding model in the table. AIC = Akaike Information Criterion; BIC = Bayesian Information Criterion.

## 4. Discussion

In the present study, 154 adolescent girls ages 10-19 years old were measured up to four times to assess the longitudinal trajectories of social motivation goals and hippocampal, mOFC, and rACC grey matter volume across age and pubertal stage. Our results demonstrated linear trajectories across pubertal development for all three social motivation dimensions (*Social Development*, *Approach Demonstration*, and *Avoidant Demonstration*), while age revealed linear changes for *Approach* and *Avoidant Demonstration Goals* but a quadratic trajectory for *Social Development Goals*. Our findings also found non-linear increases in hippocampal grey matter volume across both indices of development, and overall reductions in mOFC and rACC cortical thickness across development.

### 4.1 Social Motivation Across Adolescence

In our sample, *Social Developmental Goals* increased quadratically across age, with an inflection point around 15-years old where we observed a steeper increase. Total endorsement scores for this subdomain remained relatively elevated, reflecting an exponential increase in the prioritization of building and maintaining healthy, positive peer relationships. This can be considered a reflection of a nuanced approach to balancing both approach and avoidant demonstrative goals. Our results extend prior work that has been limited to 15- and16-year old final timepoints (Meisel et al., 2021; Trucco et al., 2014). Further, we saw linear increases in *Approach Demonstration Goals* across age and pubertal development, reflecting the increased desire to seek out positive feedback from peers. Even into later adolescence, this goal continues to steadily increase, suggesting that this is not necessarily a maladaptive behavior or cognition, when paired with the trajectories of *Social Developmental Goals* that were also observed. This could potentially reflect the increased salience and value of peer connection that girls experience across development (Pfeifer & Allen, 2012). This also algins with literature that also observes increasing approachful social goals across adolescence and even extends prior work by characterizing protracted increases into later adolescence (Meisel et al., 2021; Trucco et al., 2014).

While we found minimal change for *Avoidance Demonstration Goals* across adolescence, our LME revealed that the total endorsement of this desire remains stable and elevated during development. Like *Approach Demonstration Goals*, this may not be a maladaptive behavior but rather reflects consistent avoidance of negative peer feedback or rejection (Neel et al., 2016). When considered alongside the increases in *Social Developmental* and *Approach Demonstration Goals*, this provides more context into how adolescents navigate the prospect of negative peer evaluation while striving to create and foster positive connections. Prior work examined decreases in submissive goals (Trucco et al., 2014) across adolescence, and our results show that there is minimal change between-subjects across the developmental period. Further, this suggests that minimizing criticism or rejection from peers may be a goal that is relevant as adolescents enter young adulthood. However, despite negative peer feedback and rejection becoming more salient or meaningful across adolescence (Pfeifer & Berkman, 2018), we do not see this reflected in any increase in avoidant behaviors. Rather, our results may reflect that avoidant demonstrations are considered potentially more thoughtfully when girls make social decisions that align with *Social Developmental* or *Avoidant Demonstration Goals*.

### 4.2 Structural Neurodevelopment Across Adolescence

Leveraging LMEs, we found increased grey matter volume in bilateral HPC across age and pubertal maturation. This is consistent with our own hypothesis and previous literature (Herting et al., 2015, 2018; Vijayakumar et al., 2016). While not as robust as we were expecting, this is not as concerning as the HPC follows differentiated developmental trajectories compared to other subcortical regions (Mills & Tamnes, 2014). Further, we found protracted cortical thinning of the mOFC and rACC across development, consistent with prior literature (Goddings et al., 2019) and reflecting normative maturation and pruning of larger cortical regions of the brain. By characterizing these structural neurodevelopmental changes across age and pubertal maturation, we provide a more nuanced understanding of adolescence altogether.

Together, these findings suggest that regions implicated in social motivation continue t undergo substantial structural reorganization across adolescence. Importantly, these structural changes occurred within regions that have been implicated in valuation, affective processing, and social cognition, highlighting adolescence as a period of continued neurodevelopment in circuitry relevant to socio-affective functioning.

### 4.3 Limitations and Future Research

While our accelerated longitudinal design and integration of neurodevelopmental and behavioral data provided a strong study, several limitations warrant consideration when interpreting findings. Participant attrition across waves and incomplete cases at each wave resulted in missing data. While advanced statistical methods, including mixed-effects modeling and FIML, were employed to address this, these approaches may not fully capture individual variability.

We employed linear mixed-effects models to characterize developmental change in our variables of interest given their ability to handle complex, longitudinal data, though it might be more advantageous to use more flexible modeling strategies (i.e., generalized additive mixed models) to fully capture canonical non-linear change in adolescent neurodevelopment, and potential robustly non-linear changes in social motivational goals.

While we included two indices of adolescent development, age and pubertal maturation, we did not account for pubertal hormonal variation across maturation stages. Fluctuations in hormones such as dehydroepiandrosterone, estradiol, and testosterone are specific to phases of pubertal maturation, and may lend more precise insight into the changes observed in the current study.

The changes in our neurodevelopmental regions of interest were not directly linked to the changes observed in social motivational goals. A key concept throughout the conceptualization of this study is that HPC, mOFC, and rACC underly social motivational processes, and while this may be true, we did not run any formal statistical test or inference to verify that this is true in our sample. Further, our HPC analyses were restricted to left and right hemispheres without differentiation between anterior and posterior subregions. Emerging research indicates that longitudinal subregions of the HPC are associated different cognitive and social functions linked to motivation that may show distinct developmental trajectories (Elliott et al., 2022; Langnes et al., 2020).

Furthermore, the study’s sample demographics limit the generalizability of findings to other populations, as differences in cultural, socioeconomic, and environmental contexts may influence developmental trajectories.

### 4.4 Conclusions

This study highlights protracted developmental changes in social motivation and related structural brain maturation across adolescence. By leveraging longitudinal methods, we reveal the linear and non-linear nature of these outcomes and highlight adolescence as a crucial period for social and neural growth. Future research should build on these findings by directly examining neurodevelopmental change as it relates to social motivational changes, as well as the impact of endogenous pubertal processes including hormone secretion, variation of onset of puberty, and pubertal tempo. Further, future work should consider using more flexible modeling strategies to adequately capture canonical non-linear trajectories across our outcomes of interest. These efforts will advance our understanding of adolescent development and inform interventions that promote adaptive and positive social and emotional outcomes.

## Funding Disclosure

The project described was supported by the National Center for Advancing Translational Sciences, National Institutes of Health, through Grant Award Number TL1TR002371. The content is solely the responsibility of the authors and does not necessarily represent the official views of the NIH.

## Notes

### Competing Interest Statement

The authors have declared no competing interest.

